# Targeted delivery of RNA-based therapeutics enables functional analysis of macrophage subpopulations

**DOI:** 10.1101/2025.03.12.642907

**Authors:** Rikke Kongsgaard Rasmussen, Johann Mar Gudbergsson, Henriette Mathiesen, Line Moesgaard Strauss, Ida Holten-Møller, Mads Brejner Thomsen, Mie Wolff Kristensen, Morten Nørgaard Andersen, Anders Etzerodt

## Abstract

Macrophages infiltrate all human tissues where they play key roles in innate immunity, homeostasis, and tissue function. However, extensive clinical and experimental evidence indicates that macrophages also contribute significantly to the progression of several diseases such as cancer, cardiometabolic disorders, and inflammatory and neurodegenerative conditions. Advances in single-cell omics have revealed diverse macrophage populations in both healthy and diseased tissues. However, studying their functions is challenging due to limitations in tools for targeting specific populations. The Cre-lox system, involving Cre recombinase expression driven by macrophage-specific promoters, is widely used for gene manipulation. Despite its utility, this method has drawbacks like leaky expression, variable efficiency, and potential toxicity. Moreover, genetic models are costly and can have unintended effects on immune cells, hindering comprehensive studies on macrophage function.

To address this challenge, we developed an advanced lipid nanoparticle-based system for precise RNA therapeutic delivery to macrophages, either broadly or via antibody-mediated targeting of specific subsets. This versatile platform enables the administration of various RNA molecules, such as mRNA, siRNA, and sgRNA for CRISPR/Cas9 applications, in both *in vitro* and *in vivo* settings. It allows for targeted cell depletion or gene knockout, facilitating detailed functional analysis. Furthermore, the system’s flexibility and precision are enhanced by its compatibility with Cre-specific Cas9 expression, enabling comprehensive genomic and proteomic targeting of specific macrophage subsets.

## Introduction

Macrophages are present in all human tissues, where they play a fundamental role in health and disease^1^. These cells, now known as tissue-resident macrophages (TRMs), are seeded in the tissue early during development and are maintained locally by tissue-specific signals, either via local proliferation or replacement by long-lived monocyte-derived cells^2-4^. TRMs play an essential role in maintaining homeostasis and tissue function, and together with short-lived monocyte-derived macrophages that infiltrates inflamed tissue during infection and injury, they are recognized as a key tissue guarding cell subset that uphold tissue integrity and function^1,5^. However, extensive clinical and experimental evidence have also shown that macrophages contribute significantly to the progression of several serious pathologies such as cancer, cardiometabolic disorders, inflammatory conditions and neurodegenerative disease^1,6-9^. Macrophage heterogeneity has been recognized for several decades. However, whereas macrophage heterogeneity was earlier a matter of polarization towards pro- or anti-inflammatory phenotypes, it is now evident that TRMs and monocyte-derived macrophages are functionally and phenotypically distinct^6-8^. Within these compartments recent advancements in single-cell omics technologies have predicted that both healthy tissues and disease microenvironments are infiltrated by up to several functionally unique macrophage populations^1,10^.

Nevertheless, despite bioinformatic analyses suggesting specific roles for these subsets, obtaining direct mechanistic evidence is challenging due to the lack of easily accessible tools for targeting functional pathways in specific macrophage populations. The most common method to investigate the functional role of individual macrophage populations involves the use of macrophage specific expression of Cre recombinase to delete floxed genes or stop-codons as in the lox-stop-lox cassette to allow expression of knock-ins^11^. LysM-Cre, CSF1R and Cx3cr1 are often used to gain macrophage specific Cre expression^12^. Two main variants of the Cre system are most common; 1. Constitute active Cre, which exerts recombinase activity constantly. 2. The CreERT2 that encodes a Cre recombinase (Cre) fused to a mutant estrogen ligand-binding domain (ERT2) that requires the presence of tamoxifen for activity^13^. However, these methods are not without limitations and while the Cre-lox system is useful for conditional gene knock-out or knock-in, it can suffer from issues such as leaky expression, variable efficiency, and potential toxicity when overexpressed^14-16^. Moreover, genetic models are costly, time-consuming, and may have unintended effects on certain immune cell populations, such as the adverse impact of tamoxifen on peritoneal macrophages^17,18^ prohibiting widespread screening effort to properly understand function of individual macrophage populations.

To address this challenge, we have developed a new lipid nanoparticle (LNP)-based toolbox for targeted delivery of RNA therapeutics to macrophages in general or via antibody-mediated targeting to individual subsets. This system allows delivery of various RNA molecules, such as messenger RNAs (mRNAs), small interfering RNA (siRNA) and single guide RNAs (sgRNAs) for CRISPR/Cas9 *in vitro* and *in vivo*, enabling the depletion of cells or knockdown or knockout of specific genes to facilitate functional analysis. The platform offers a multitude of flexibility and precision and can be combined with Cre-specific expression of Cas9 enzymes, enabling both genomic and proteomic targeting of individual macrophage populations.

## Methods

### LNP formulation

RNA-loaded LNPs were prepared using the NanoAssemblr® Ignite platform using the lipid formulation specified in table 1. In brief, RNA in the aqueous phase (100 μM Sodium Acetate buffer, pH 4.5, Thermo Scientific) was mixed with the lipid phase in a flow rate ratio of 4:1 (aq:lipid), with a total flow rate of 5ml/min. After formulation, the LNPs were dialyzed twice against 1X PBS (Cytiva) in 14 kDa MWCO dialysis tubes (Spectra/Por® 4) overnight and concentrated by covering dialysis tubing with PEG (Mw 35,000) (Sigma-Aldrich) for 10-20 min at room temperature. Following, LNPs were sterile filtered using cellulose Acetate 0.2 μm membrane filter (Avantec®) and LNP-lipid concentration was analyzed using high-performance liquid chromatography on a Dionex Ultimate3000 HPLC system (Thermo Fisher Scientific) equipped with a Nucleosil 100-3C18 column. RNA concentration was determined using the modified Quant-iT RiboGreen RNA (Thermo Scientific #R11490) protocol proposed by Precision NanoSystems. To ensure long-term stability of the RNA, LNPs were stored in −70°C, using 5% sucrose (Merck) as a cryoprotectant ^19^.

**Table 1.** LNP formulation table.

Abbreviations: DSPC – Distearoylphosphatidylcholine. DMG-PEG(2000) - 1,2-dimyristoyl-rac-glycero-3-methoxypolyethylene glycol-2000. DSPE-PEG(5,000)-DBCO - 1,2-distearoyl-sn-glycero-3-phosphoethanolamine-n-[dibenzocyclooctyl(polyethylene glycol)-5000].
|  | Lipid ratio<br>Untargeted LNPs | Lipid ratio<br>Targeted LNPs | Manufacturer |
| --- | --- | --- | --- |
| SM-102 | 50 | 50 | BOC Sciences, |
| DSPC | 10 | 9 | Cayman |
| Cholesterol | 39 | 38 | Sigma Aldrich |
| DMG-PEG(2,000) | 1 | 5 | Cayman |
| DSPE-PEG(5,000)-DBCO | 0 | 0.5 | Avanti Polar Lipids |

### RNA molecules

mRNA was either purchased from Trilink (mCherry) or prepared using the Takara IVTpro™ mRNA synthesis kit, combined with CleanCap reagent AG (TriLink), according to the manufacturer instructions. In all formulations uridine was substituted with 5-mothoxyuridine (Jena Bioscience). For mRNA templates, constructs were cloned into an IVT plasmid (pUC57, Addgene) that contained previously described 5’ and 3’ untranslated regions with a 120 nt poly(A) tail^20-23^, as well as the T7 promotor and the required AG site for 5’ capping. Sequence can be obtained on request. siRNA was purchased from Thermo Fischer Scientific, using their SilencerSelect RNA (table 2). sgRNA were designed using the CCTOP’s online design tool ^24,25^ (table 3), and ordered from IDT.

**Table 2.** siRNA table.

| Target | ID |
| --- | --- |
| mCD9 | n523179, n523180, s201141 |
| hCD9 | s2597, s2598 |
| mLyve1 | 100109 and 100110 |
| Negative control | 4390844 |

**Table 3.**
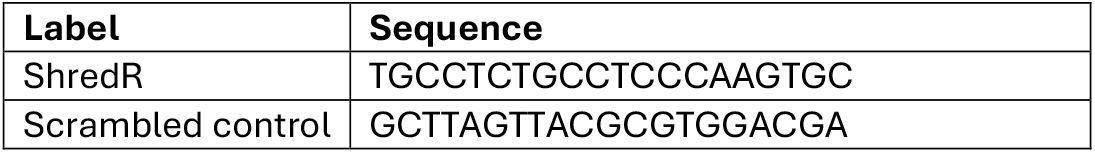
sgRNA table.

### Antibody conjugation to LNPs

InVivoMAb Anti-mouse TIM4 antibody (Bio X Cell) was functionalized in a 2 step process. Firstly, the antibody was oxidized using 10 mM sodium periodate (Thermo Fischer Scientific), 150 mM NaCl (Merck), and 100 mM sodium acetate for 20 minutes at room temperature. 10X PBS (Cytiva) was added to achieve 1X PBS in order to stop the reaction, and excess sodium periodate was washed away. Following oxidation, a functional azide group was attached; Oxidized antibody was mixed with Aminooxy-PEG(3)-Azide (Conju-Probe), 1X aniline catalyst (Biotium), 150 mM NaCl, and 100 mM sodium acetate for 3 hours at room temperature. Again, 10X PBS was added to achieve 1X PBS in order to stop the reaction, and unbound aminooxy-PEG(3)-Azide and aniline catalyst was washed away, and buffer exchanged to 1X PBS.

LNPs and functionalized antibody were mixed in a ratio of 1:2500, approximate antibody concentration of 0.5 mg/ml during conjugation. The mix was left to incubate over the weekend at 4°C.

### Bone marrow derived macrophages (BMDMs)

Murine BMDMs were prepared by flushing the femur and tibia of C57/Bl6 female mice age 7-12 weeks with RPMI-1640 (Gibco). Following, bone-marrow was filtered through a 70 μm cell strainer (pluriSelect) and 10 × 10^6^ bone-marrow cells were plated in uncoated petri dish (Greiner Bio-one) in RPMI-1640 with 10% FBS (Biowest), 100 U/ml penicillin, 100 μg/ml streptomycin (Gibco), 2 mM L-Glutamine (Gibco) and 25 ng/ml m-CSF (Peprotech). Cells were supplemented on day 3 with the same media as above and plated for experiments on day 6.

Lipofectamine 2000 (Invitrogen) was used as an alternative to LNP based RNA delivery *in vitro*, according to the manufacturer’s instructions. Lipofectamine, RNA and LNPs were dissolved in Opti-MEM (Gibco) and no pen/strep was used in the media during the experiment. Cells were harvested for flow cytometry using 1X PBS (Cytiva) with 10 mM EDTA (Invitrogen), until visually confirmed that the cells had detached from the plate. Cells were live/dead stained using SYTOX Blue (Invitrogen) and samples were run on NovoCyte Quanteon 4025 flow cytometer equipped with four lasers (405 nm, 488 nm, 561 nm and 637 nm) and 25 fluorescence detectors (Agilent), using the NovoExpress software (v. 1.6.2, Agilent). Analysis was performed in FlowJo (v. 10.10.0).

### Human monocyte derived macrophages (MDMs)

Human peripheral blood mononuclear cells (PBMCs) were purified from buffy coats obtained from the blood bank at the Department of Clinical Immunology, Aarhus University Hospital, Aarhus, Denmark. Buffy coats were diluted 1:2 in 0.9% NaCl, and the PBMCs were isolated using density gradient centrifugation (400 g, 30 min, room temperature) on a Histopaque-1077 gradient (Sigma-Aldrich). The PBMCs were washed twice in PBS with 2% fetal bovine serum (FBS; Sigma-Aldrich) and 1 mM EDTA (Merck Millipore) before monocytes were isolated by negative selection using the EasySepTM Human Monocyte Isolation Kit (StemCell Technologies) according to the manufacturer’s instructions. After isolation and wash, the monocytes were cultured for 7 days in RPMI-1640 medium (Biowest) with 10% heat-inactivated FBS, 100 U/ml penicillin, 100 μg/ml streptomycin, and 2 mM gluta-mine (Gibco) (complete RPMI-1640). For MDM differentiation, the medium was further supplemented with 10 ng/ml M-CSF (day 1-5), 1 ng/ml GM-CSF (day 1-5), and 10 ng/ml IL-10 (day 5-7) (all from Pepro-Tech).

### Mice

All animals were housed in the animal facility at the Department of Biomedicine, Aarhus University. Water and food was provided ad libitum and with a 12 hour night/daylight cycle. C57/Bl6 female mice, age 7-12 were ordered from Janvier. The *in vivo* ShredR model was performed using C57/Bl6-H11^Cas9^, strain 028239, bred in-house, generously gifted from Martin Thomsen, Department of Biomedicine, AU. CD115-Cre x Cas9-EGFP^fl/fl^ mice used for BMDM viability assay, were prepared by breeding CD115-Cre and Cas9-EGFP^flfl^, which were a generous gift from Felicity Davies and Joanna Kalucka, Department of Biomedicine, AU, respectively.

### *In vivo* experiments

Mice were injected i.p. with the indicated LNPs, and after 24-48 hours, peritoneal lavage was harvested for flowcytometric analysis. Peritoneal lavage was obtained by injecting 5 ml 1X PBS + 2mM EDTA into the peritoneal cavity of mice, after cull. The mice were then quickly massaged on the stomach, and the peritoneal wash fluid was recovered. The peritoneal wash was counted, and 1 × 10^6^ cells were resuspended in Fc blocking buffer (FACS buffer (1X PBS + 2% FBS + 2mM EDTA + 0.09% sodium azide) with αCD16/32-clone 24G2) for 15 minutes at 4°C. The cells were then stained with antibodies in the same blocking buffer with additional OligoBlock ^26^ and Brilliant Stain buffer (BD Horizon) for 30 minutes at 4°C. The cells were washed and resuspended in FACS buffer. Immediately prior to running the samples, SYTOX dyes (Invitrogen) were added to stain dead cells. Flow cytometry was run on a five lasers (355 nm, 405 nm, 488 nm, 561 nm and 637 nm) ID7000™ Spectral Cell Analyser (SONY Biotechnologies) or a Novocyte Quanteon flow cytometer. Samples run on the ID7000 were recorded and analyzed using the ID7000 Acquisition and Analysis software version 2.0.2 (Sony Biotechnologies) utilizing the Weighted Least Squares Method (WLSM) algorithm for spectral unmixing. Final analysis for both Quanteon and ID7000 data was analyzed in FlowJo v. 10.10.0.

**Table 4.** Antibodies for flow cytometry.

|  | Clone | Fluorochrome | Manufacturer # |
| --- | --- | --- | --- |
| NK1.1 | PK136 | BUV395 | BD Horizon<br>#564144 |
| CD90.2 | 30-H12 | BUV496 | BD Optibuild<br>#741047 |
| CD5 | 53-7.3 | BUV563 | BD Optibuild<br>#741215 |
| CD102 | 3C4(mIC2/4) | BUV615 | BD Optibuild<br>#751392 |
| CD11b | M1/70 | BUV661 | BD Horizon<br>#612977 |
| Tim4 | RMT4-54 | BUV737 | BD Optibuild<br>#749134 |
| CD19 | 1D3 | BUV805 | BD Horizon<br>#568287 |
| SYTOX-Blue | N/A | N/A | Invitrogen<br>#S34862 |
| CD163 | TNKUPJ | eFluor450 | eBioscience<br>#48-1631-82 |
| Ly6G | 1A8 | BV570 | Biolegend<br>#127629 |
| Ly6C | AL-21 | BV605 | BD Horizon<br>#563011 |
| F4/80 | T45-2342 | BV650 | BD Optibuild<br>#743282 |
| CD115 | AFS98 | BV711 | Biolegend<br>#135515 |
| CD163 | TNKUPJ | SB702 | eBioscience<br>#67-1631-82 |
| SiglecF | E50-2440 | BV786 | BD Optibuild<br>#740956 |
| CD45 | 30-F11 | AF532 | Invitrogen<br>#58-0451-82 |
| CD11c | N418 | BB700 | BD Optibuild<br>#745852 |
| CD64 | X54-4/7.1 | PerCP-Cy5.5 | Biolegend<br>#139308 |
| Lyve1 | ALY7 | PE | eBioscience<br>#12-0443-82 |
| CD11b | M1/70 | PE-Cy5 | eBioscience<br>#15-0112-81 |
| CD209b | REA125 | PE-Vio770 | Miltenyi Biotec<br>#130-106-332 |
| CD226 | 10E5 | PE-Cy7 | BD Pharmingen<br>#567651 |
| CD169 | Ser4 | eFluor660 | eBioscience<br>#50-5755-82 |
| MHC-II | M5/114.15.2 | AF700 | eBioscience<br>#56-2321-80 |
| Ly6G | 1A8 | APC-Cy7 | BD Pharmingen<br>#560600 |

### Incucyte

Adherent human MDMs were harvested by 15 min incubation with PBS/0.5% bovine serum albumin (BSA) (Merck Millipore)/5 mM EDTA/4 mg/ml lidocaine (Sigma-Aldrich) followed by detachment using a cell scraper. The MDMs were seeded in a 96 well plate (Sarstedt) (5000 cells/well in complete RPMI) and left in the incubator for a few hours before addition of LNPs containing mRNA for GFP (TriLink Biotechnologies). The uptake of LNPs and subsequent release and translation of functional mRNA was monitored and quantified over time by measuring the GFP signal in the green fluorescence channel on the Incucyte® S3 Live-Cell Analysis System. The plate was scanned every 1 h for 3 days with 2 images/well using a 20× objective and an acquisition time of 300 ms. Each condition was run in duplicate. The GFP signal (total integrated green fluorescence intensity) was calculated using the Incucyte 2022B Rev2 software.

### qPCR

qPCR gene expression assays were performed using the KAPA SYBR® fast low ROX (Sigma Aldrich, #KK4620), 5 ng RNA, and 0.4 μM combined forward/reverse primer. Assays were then run on a LightCycler® 480 System (Roche). ΔCT values were obtained by normalizing gene expression to *ppia* or *PPIB*.

**Table 5.**
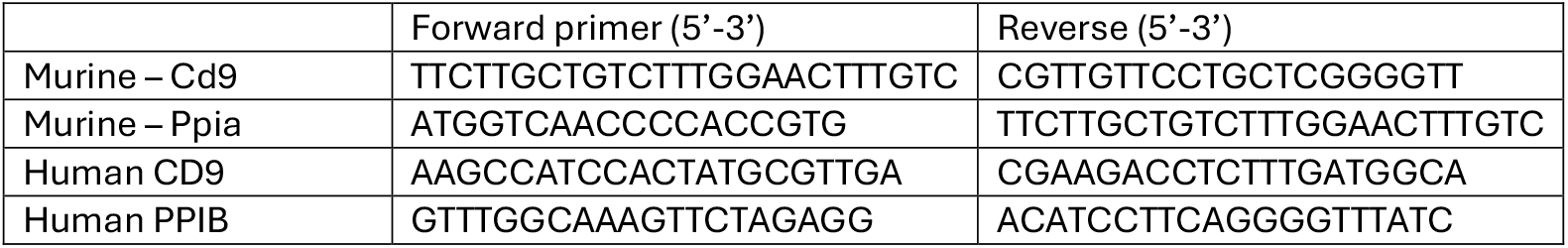
qPCR primers.

### Viability assay

BMDMs were prepared as described above, from CD115-Cre x Cas9-EGFP^fl/fl^ mice. On day 6, the BMDMs were plated in uncoated 24 well plates (Falcon, #10396881) with the indicated dose of RNA-LNPs. Cells were harvested by collecting the supernatant and pooling with any adherent cells, harvested using 1X PBS with 10 mM EDTA. Viability and cell count was assessed using the Nucleocounter NC-3000™ with the Nucleoview NC-3000 version 2.2.0.0 (Chemometec) and Solution 13 (Chemometec).

### Sorting of peritoneal cavity cells

Peritoneal cavity cells were harvested from 2 naïve 11-week-old C57/Bl6 mice, and then counted and multiplexed using the hashtag-oligo antibodies (TotalSeq, Biolegend). The cells were mixed in a 1:1 ratio and stained for sorting using the antibodies in the table below, mixed with Fc blocking buffer described previously. Staining was performed for 30 minutes at 4°C in the dark. After staining, the cells were washed and resuspended in FACS buffer with viability dye. Immediately prior to sorting, pleuronic acid (Thermo Scientific) was added to a final concentration of 0.3% (w/v)

**Table 6.**
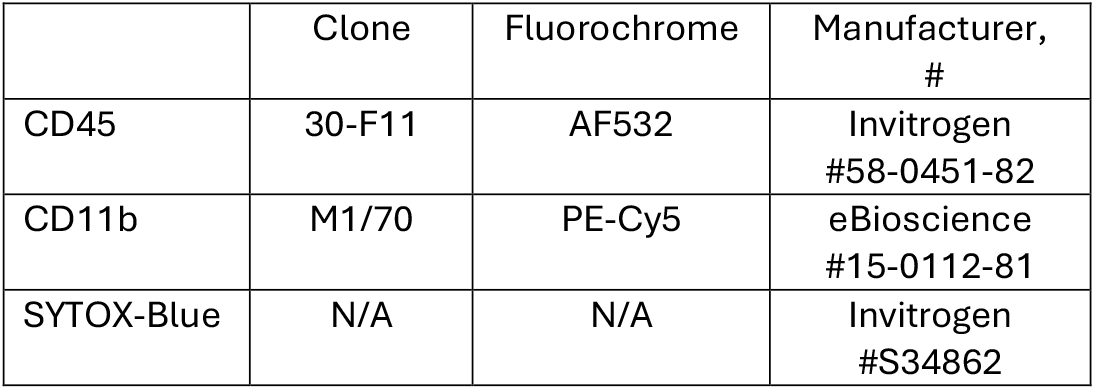
Antibodies and dyes used for FACS.

### Single cell RNA sequencing (scRNA-seq) and analysis

scRNA-seq was completed using the ASTERIA Benchtop 3’ scRNA-Seq Sample preparation kit (Scipio Bioscience), according to the manufacturers protocol using 30,000 cells as input. cDNA was amplified with the addition of HTO additive primer v2 (see table). Sample cleanup was executed by using SPRIselect (Beckman Coulter), and HTO and mRNA were separated to generate separate libraries. The quality of the cDNA was assessed using the Bioanalyzer 2100 (Agilent), using their high sensitivity DNA kit (Agilent).

Library prep was performed using the NEBNext® Ultra™ II DNA Library Prep Kit according to the manufacturers protocol, except a few modifications to ensure concordance with the ASTERIA protocol. This involved exchanging the Universal PCR primer/i5 with the Asteria Library prep primer. PCR was run as recommended, as was cleanup, and the quality of the library was assessed using the Bioanalyzer 2100, with the High Sensitivity DNA Kit. HTO library was prepared according to BioLegend’s TotalSeq HTO library preparation protocol, with KAPA HiFi HotStart ReadyMix (Roche). Cleanup was performed using SPRIselect. Subsequently, libraries were sequenced to a depth of on average 100,000 reads per cell using a NovaSeq S1 flow cell (Illumina) averaging to approximately 2000 genes per cell.

**Table 7.**
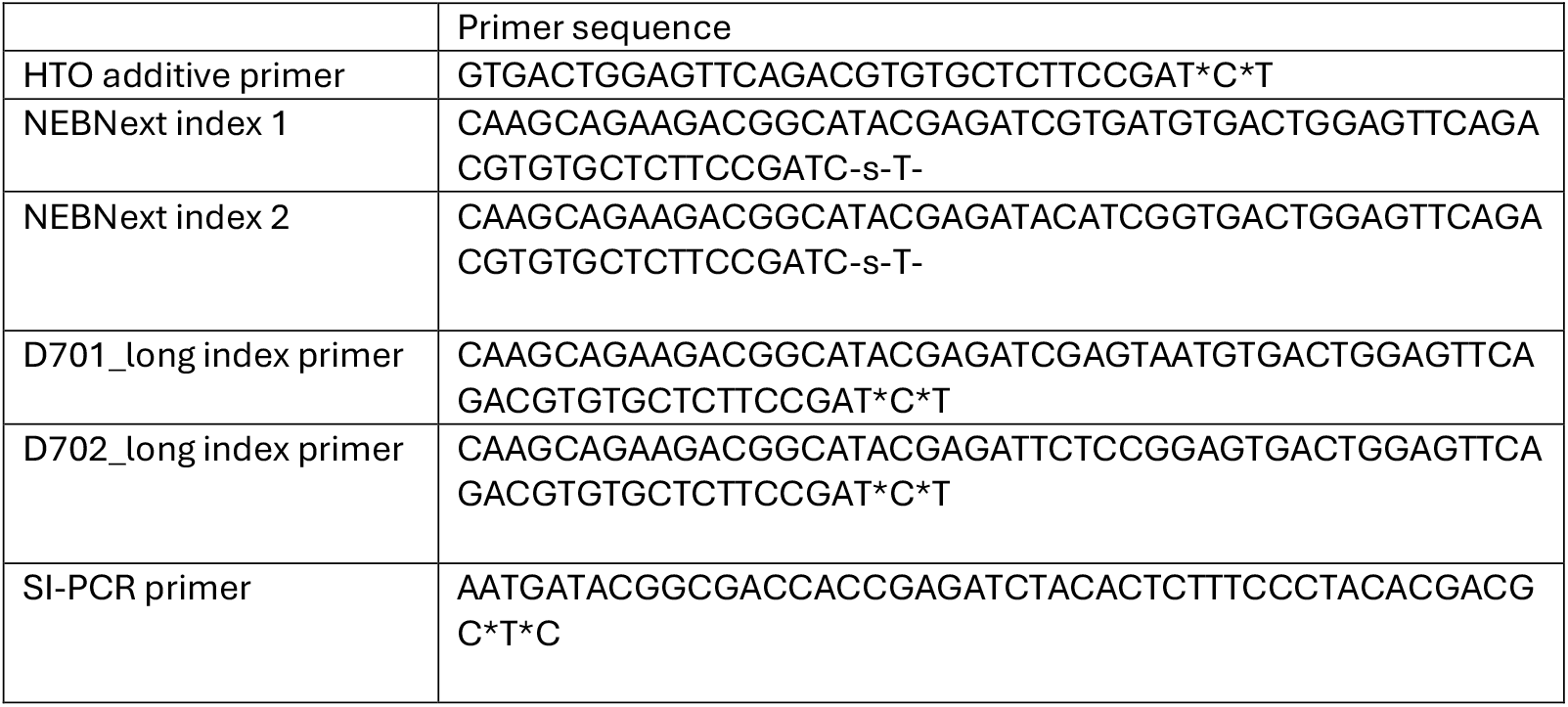
Primers for scRNA sequencing.

After sequencing, samples were processed with the Cytonaut platform from Scipio Bioscience (v7) using Mouse_Scipio_2023_A reference genome (source: ensemble, version: GRCm38). Filtered read count matrices were analyzed with Seurat (v5.1.0) running in R (v4.2.0). Samples were filtered to remove cells with high mitochondrial read contamination (>2%), low gene detection rate (<1000) and high gene detection rate (>5000) or high read count (>20000). After filtering, 17574 genes and 3094 cells remained. Cells were normalized and scaled using default settings in Seurat. Cell type annotation was done using the single-cell Mouse Cell Atlas (scMCA v0.2.0) and cells annotated as either monocytes or macrophages were kept resulting in a total of 3080 cells. The 2000 most variable genes were used to calculate the principal components. Cells were clustered with the Louvain algorithm using 9 principal components and a resolution of 0.9. Differentially expressed genes were calculated by comparing each cluster to all other clusters using FindAllMarkers(logfc.threshold = 0.25, min.pct = 0.10, only.pos = TRUE) with default Wilcoxon Rank Sum test. The top 10 markers were selected by first sorting by the lowest P-adjusted value and subsequently by the highest log2 fold-change.

### Statistics

Statistical tests were performed when appropriate, and the statistical test is indicated in the figure legend. Two-way ANOVA with multiple comparisons was used with Turkeys correction. The student’s t test was used for single column separated data, Dunnett’s test for multiple columns. All statistical tests were performed in Prism, version 10.3.1. Error bars indicate ±SEM, **** P<0.0001, *** p<0.001, ** p<0.01 *p<0.5.

## Results

### Lipid nanoparticles efficiently deliver RNA to primary macrophages *in vitro*

To ensure effective delivery of functional RNA to macrophages, we encapsulated RNA in LNPs using a lipid formulation similar to the Spikevax COVID-19 vaccine. This includes the commercially available ionizable lipid SM-102, DMG-PEG, cholesterol and DSPC (fig. 1A). Our aim was to develop a versatile formulation capable of efficiently encapsulating both long and short RNA molecules, such as mRNAs, siRNA and sgRNAs. Utilizing this formulation, we achieved robust and efficient loading, with high mean loading efficiencies across the different RNA types of 93.8% (mRNA), 95.4% (siRNA) and 93.9% (sgRNA) (fig. 1B).

**Figure 1.**
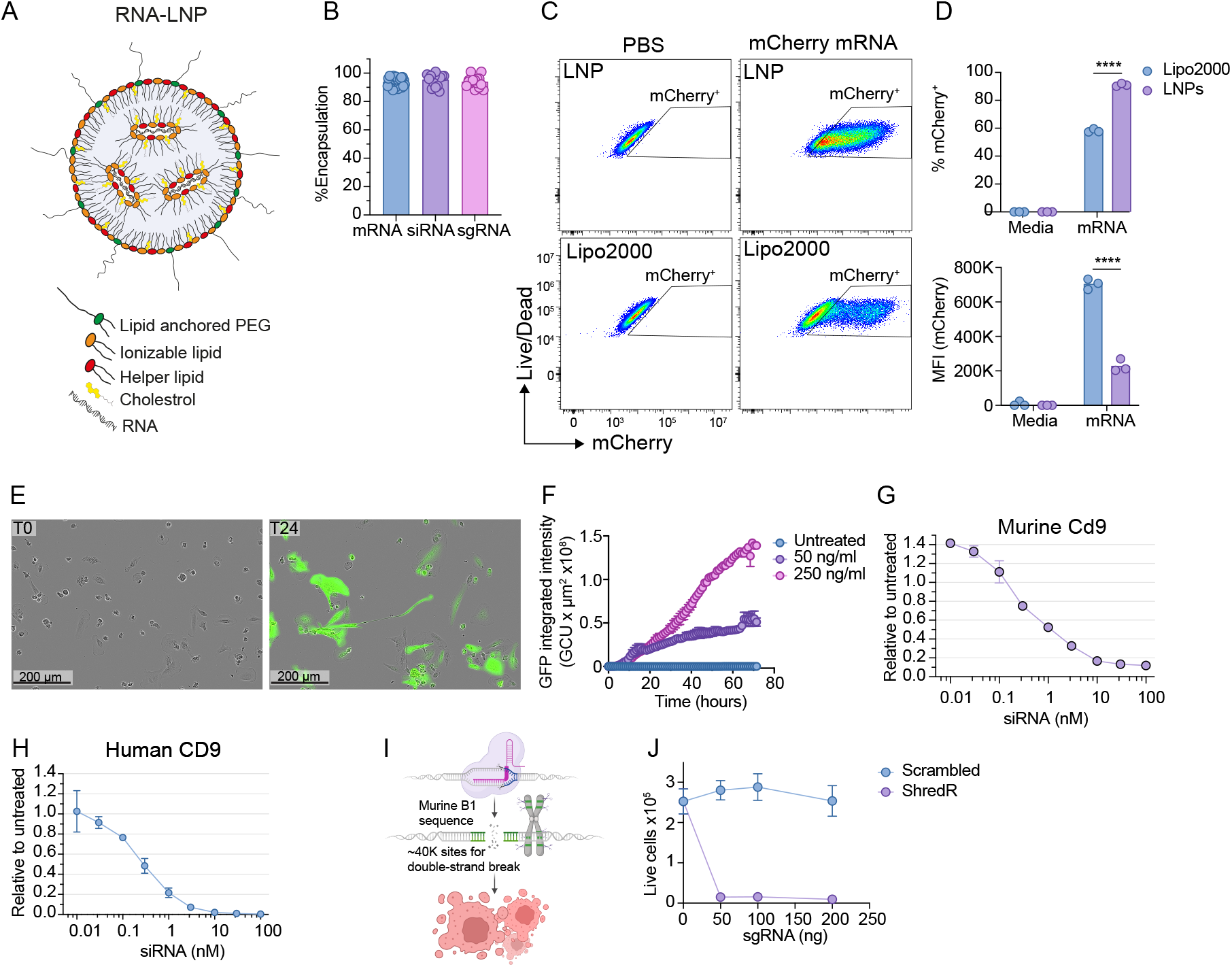
In vitro characterization of SM-102 LNPs. (A) schematic of LNP formulation comprising of 4 major lipid components encapsulating RNA. (B) Encapsulation efficiency of various RNA types. (C) Comparison of lipofectamine and LNP mediated delivery of mCherry mRNA in murine BMDMs after stimulation 48 hours. (D) Quantification of percent of mCherry positive cells and mCherry MFI. (E) Fluorescent imaging of eGFP expression in human monocyte derived macrophages in the same well, 0 hours and 24 hours after stimulation with LNP encapsulating eGFP mRNA. (F) Quantification of the eGFP intensity over time. Titration of Cd9 siRNA in bone marrow derived macrophages using siRNA directed against murine Cd9 (G) or in human monocyte-derived macrophages using siRNA directed against human CD9 (H). (I) Schematic of the ShredR strategy introducing double stranded breaks in an estimated 40.000 sites in the genome, resulting in cell death. (J) Cell count and viability of Cas9 expressing bone-marrow derived macrophages after 72 hours stimulation with LNPs encapsulating either ShredR or Scrambled sgRNA. Error bars show ± SEM and for (D) statistically significant differences was calculated using 2-way ANOVA followed by Turkeys multiple comparison test, ****, P < 0.0001.

Macrophages are notoriously difficult to deliver nucleic acids to, primarily due to their biological characteristics and specialized functions^27^. To explore the capability of the SM-102 formulated LNPs to mediate RNA delivery to macrophages, we initially compared the transfection efficiency of mRNA formulated LNPs with a commonly used transfectant (Lipofectamine2000) in murine BMDMs using mCherry mRNA as reporter. After incubation for 24 hours, flow cytometric analysis revealed that SM-102 formulated LNPs efficiently delivered RNA to most BMDMs, with an average of 91% of the BMDMs expressing mCherry (fig. 1D). In contrast, transfection with Lipofectamine2000 resulted in a much lower transfection efficiency with only 58% of BMDMs becoming mCherry positive, although the MFI of the reporter in the mCherry+ gate was higher compared to LNPs. In addition to a lower transfection efficiency, treatment with lipofectamine2000 also had a markedly negative effect on macrophage viability compared to LNPs and media control (fig. S1B).

Next, we sought to evaluate the ability of the SM-102 LNPs to deliver RNA to primary human monocyte derived macrophages (MDMs). This was achieved using live imaging microscopy and GFP mRNA as reporter. Upon incubation, the SM-102 LNPs were rapidly endocytosed by the MDMs, with GFP expression detectable as early as 6 hours after start of incubation (fig. 1E-F). The GFP signal intensity increased over time as the LNPs continued to be internalized, with minimal impact on MDM morphology (Fig. 1E).

Following evaluation of transfection efficiency of the LNPs, we next aimed to evaluate the ability of SM-102 LNPs to efficiently deliver siRNA to both murine and human primary macrophages. Using siRNA directed against CD9 as an example, we observed an efficient and dose-dependent knock down of CD9 resulting in a significant reduction of CD9 mRNA expression both in murine macrophages (17% expression of controls) and human macrophages (2 % expression of controls) (fig. 1G-H).

In addition to mRNA and siRNA, LNPs also allow delivery of components of the CRISPR/Cas9 gene editing system that allows for a wide variety of modulating of gene expression such as knockout, knock-in, and temporary up- or down-regulation of specific target genes via CRISPR activation (CRISPRa) or inhibition (CRISPRi) techniques. Moreover, Glow and Colleagues recently showed that the CRISPR/Cas9 system may also be used for efficiently depletion of human cells by targeting specific short interspersed nuclear elements (SINEs) that are distributed throughout the genome in high numbers^28^. SINEs are also present in the murine genome and to develop an efficient method for depletion of macrophage subsets using RNA-loaded LNPs, we designed new sgRNA targeting SINEs in mouse based on the principles described by Glow et al. The resulting sgRNA (ShredR, Fig. 1I) was subsequently encapsulated in SM-102 LNPs and incubated with Cas9-EGFP-BMDMs. ShredR loaded LNPs efficiently depleted all BMDMs as cell counts after 72 hours incubation was reduced below the detection limit, whereas BMDMs incubated with LNPs encapsulating scrambled control were unaffected (fig. 1J).

In summary, these results show that SM-102 formulated LNPs allows for a highly efficient encapsulation of RNA and delivery to both murine and human primary macrophages *in vitro*.

### LNPs deliver RNA mainly to macrophages in the peritoneal compartment

Next, we sought to investigate if the SM-102 formulated LNP would also allow for efficient delivery of RNA to macrophage *in vivo*. To this end, mice were injected *i*.*p*. with 100 ng mCherry mRNA encapsulated in SM-102 LNPs. After 24 hours, peritoneal cells were harvested by peritoneal lavage and mCherry expression was analyzed by flow cytometry (gating strategy available in fig. S2A-B). Interestingly, not only were the SM-102 LNPs able to deliver mRNA to peritoneal cavity macrophage *in vivo*, but macrophages were also the primary cell type taking up the LNPs, with an average of 91.4% of macrophages being mCherry positive. In addition to macrophages, 13.6% of monocytes were targeted by the SM-102 LNPs (fig. 2C) whereas minimal or no uptake was observed in the remaining peritoneal cells (fig. 2D).

**Figure 2.**
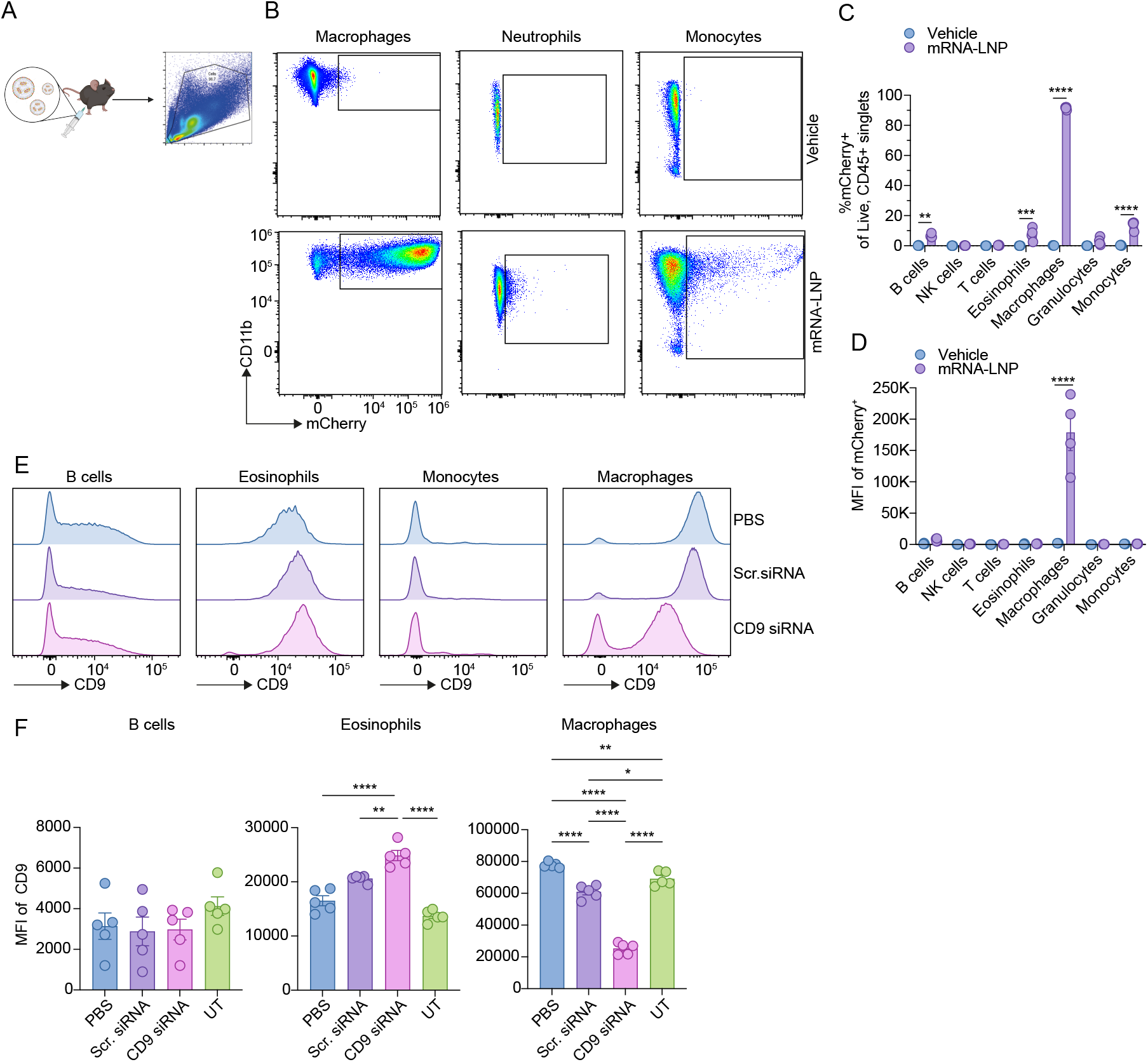
in vivo uptake of SM102 LNPs is mainly restricted to macrophages. (A) Experimental layout illustrating i.p. injection of SM102 LNPs encapsulating mCherry mRNA in C57BL/6j mice followed by spectral flow cytometric analysis of the peritoneal immune compartment after 24 hours. (B) Flow cytometric analysis of mCherry expression in monocytes, neutrophils and macrophages in vehicle injected (top) and mRNA(mCherry)-LNPs (bottom) injected mice. (C) % mCherry positive cells out of the total immune cell compartment and (D) mCherry MFI of the mCherry positive cells. (E) Histograms showing flow cytometric analysis of CD9 expression in B cells, eosinophils, monocytes and macrophages harvested from peritoneum after treatment with LNPs encapsulating siRNA against mouse CD9 or controls and (F) MFI of CD9 on B cells, eosinophils and macrophages. Error bars show ±SEM and statistically significant differences was calculated using 2-way ANOVA followed by Turkeys multiple comparison test, **, P < 0. 01, ***, P < 0.001, ****, P < 0.0001

To further explore if SM-102 LNPs also allow for delivery of relevant levels of functional RNA such as siRNA, we next injected with mice with in vitro functional SM-102 LNPs containing CD9 siRNA 3 times every 48 hours. 72 hours following the final injection, peritoneal lavage was harvested and subsequent flow cytometric analysis of CD9 expression on peritoneal immune cells showed a specific and prominent CD9 KD in macrophages (fig. 2E, F), with no effect in total cell number or in the frequency of macrophages in the peritoneal compartment (sup. fig 2), indicating tolerance of repeated injections.

### Macrophages are phenotypically heterogenous, and represent subpopulations identified at single cell and protein level

To evaluate the potential of designing SM-102 LNPs for targeting of distinct macrophage subsets, we next evaluated macrophage heterogeneity in the peritoneal cavity by performing single cell transcriptomic analysis of macrophages isolated from naïve mice. Macrophages were isolated by flow assisted cell sorting, gating on CD45^+^ CD11b^+^ cells and processed for scRNA-seq analysis using the SciPio Asteria scRNAseq benchtop kit. As expected, the analysis revealed considerable heterogeneity within the macrophages compartment, predicting 10 different clusters of peritoneal macrophages (PCM, fig. 3A, B). Extensive work has previously defined two separate peritoneal macrophage populations. These are the GATA6 dependent and tissue resident large peritoneal macrophages (LPMs) and the monocyte-derived and Irf4 dependent small peritoneal macrophages (SPMs). When overlaying the specific signature genes of LPMs (*Gata6, Timd4, Icam2 and Adgre1*) and SPMs (*Irf4, Cd226, MHC II*) on the predicted clusters, we observed 5 clusters (PCM_1-4,6,7) of LPMs, a single cluster of SPMs (PCM_9), and three clusters of peritoneal macrophages that had expression of both LPM and SPM signature genes (PCM_5,8,10). These clusters were identified as intermediate peritoneal macrophages (intPM, fig. 3C).

**Figure 3.**
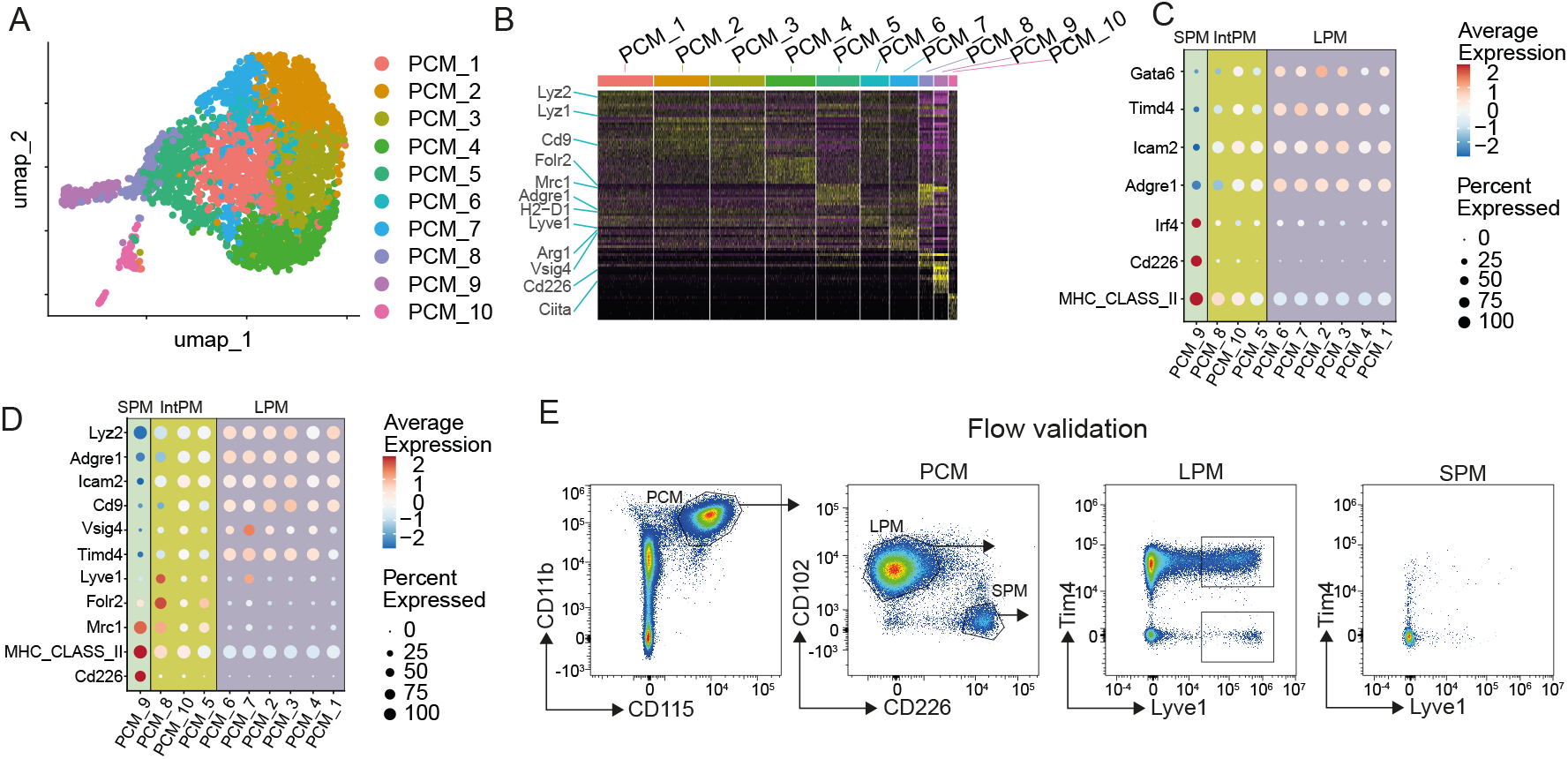
Macrophage heterogeneity in the peritoneal compartment reveals several subpopulations. (A) Unsupervised dimensional reduction and clustering of scRNA-seq data by UMAP from peritoneal cavity macrophages isolated by flow assisted cell sorting on CD45+, CD11b+ cells from peritoneal lavage of 2 Naive C57BL/6j mice. (B) Heatmap of the top 10 differentially expressed genes in each cluster and (C) dot plot of gene expression of signature genes from small peritoneal macrophages (Irf4, Cd226, MHCII signature) and large peritoneal macrophages (Gata6, Timd4, Icam2, Adgre1) in, small peritoneal macrophages (SPM), intermediary peritoneal macrophages (intPM) and large peritoneal macrophages (LPMs). (D) Dot plot of the expression of macrophage markers previously used in flow cytometry and (E) validation of Lyve1 expression of both Tim4+ and Tim4-peritoneal macrophages using spectral flow cytometry.

Interestingly, when further exploring expression of the differentially expressed cluster markers that are also used as markers in flow cytometry (Fig. 3B,D), we observed that, whereas most markers, including *Adgre1* (F4/80), *Icam2* (CD102), *Timd4* and *Cd226* only allows for segregation of populations in LPM and SPM, *Lyve1* clearly defined a single population of LPM (PCM_7) and intPM (PCM_8). To validate this observation, we next performed flow cytometric analysis of Lyve1 expression on the peritoneal cavity macrophage compartment. Macrophages were first gated as CD11b^+^ CD115^+^ and subsequently separated in SPM and LPM by gating on CD102^+^, CD226^-^ (LPMs) and CD102^-^, CD226^+^ (SPM) cells (fig. 3E). Next, we analyzed Tim4 and Lyve1 expression, and found, as predicted by scRNAseq analysis, two separate populations within the CD102^+^ macrophage gate, namely the Tim4^+^, Lyve1^+^ population that corresponds to PCM_7 cluster and the Tim4^-^, Lyve1^+^ population that corresponds to the PCM_8 cluster. SPMs were mainly Tim4^-^, Lyve1^-^.

### Targeted LNPs deliver RNA to subpopulations of macrophages

Having successfully identified multiple clusters of peritoneal macrophages, we next set out to modify the formulation our SM-102 LNPs to allow specific delivery of RNA to individual populations (fig. 4A). This included increased concentration of poly-ethylene glycol (PEG) containing lipids to 5% of total lipid concentration to reduce general uptake and increase circulation time as previously described^29-31^. Furthermore, we included a click-chemistry functionalized PEGylated lipid, enabling attachment of antibodies using copper-free click chemistry. To conjugate antibodies to LNPs, the antibodies were functionalized with azide-groups and covalently attached to DBCO available at the distal end of lipid-PEG in the outer LNP membrane, similar to previously published methods^32^.

**Figure 4.**
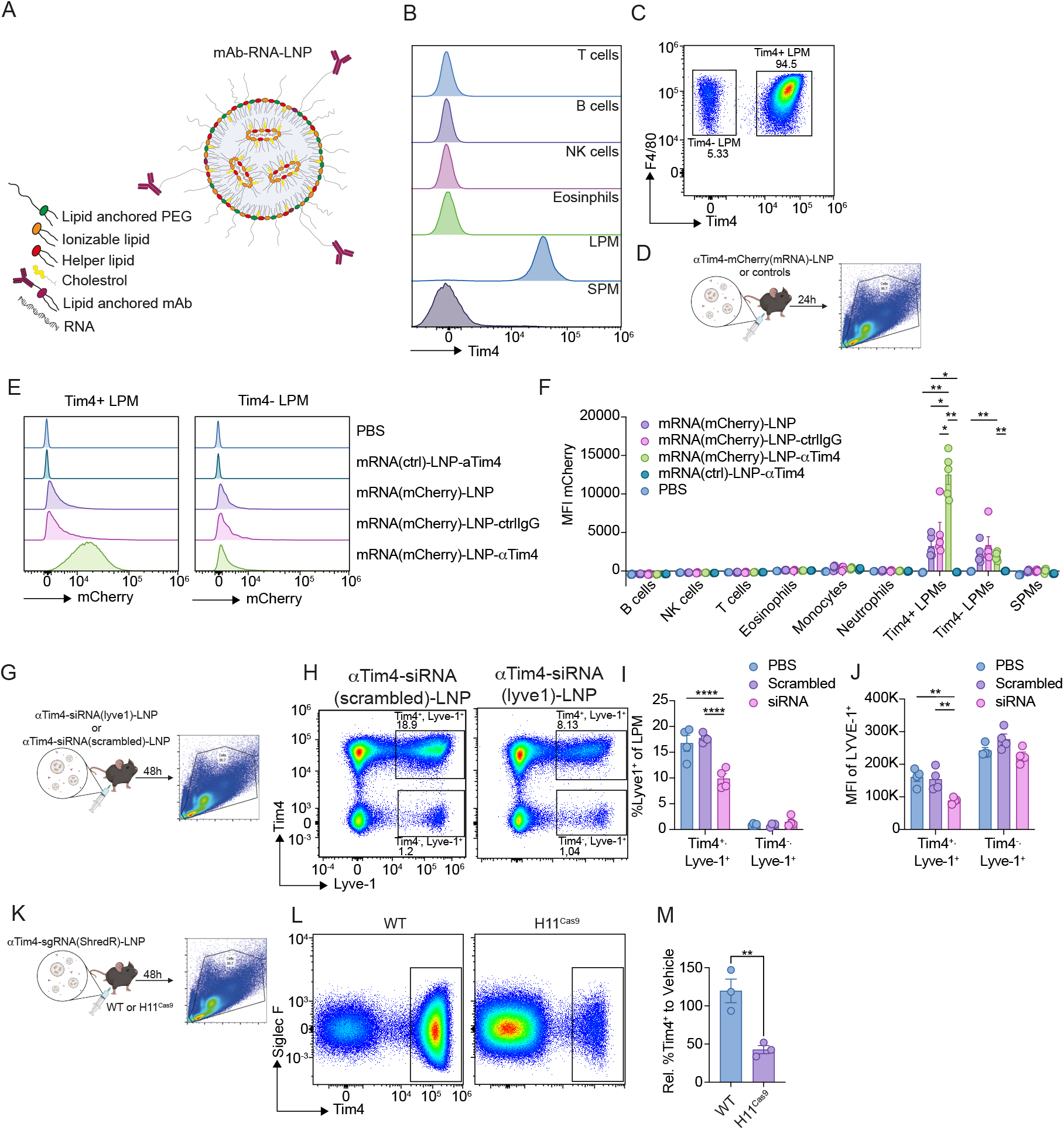
In vivo characterization of targeted SM-102 LNPs. (A) schematic overview of LNPs-conjugated covalently to antibodies using copper-free click chemistry. (B) Histrogram showing expression analysis of Tim4 on immune cells harvested from the peritoneal cavity from a naive mouse and (C) gating of Tim4+ and Tim4-LPMs. (D) Experimental layout illustrating i.p. injection of SM102 LNPs encapsulating mCherry mRNA conjugated with anti-mouse Tim4 mAb (mRNA(mCherry)-LNP-αTim4) or controls in C57BL/6j mice followed by spectral flow cytometric analysis of the peritoneal immune compartment after 24 hours. Controls include anti-mouse Tim4 mAb conjugated LNP encapsulating non-mCherry mRNA (mRNA(ctrl)-LNP-αTim4), LNP encasulating mCherry mRNA conjugated to isotype ctrl IgG (mRNA(mCherry)-LNP-ctrlIgG), non-conjugated LNP encasulating mCherry mRNA (mRNA(mCherry)-LNP) or PBS. (E) Flow cytometric analysis of mCherry expression in LPM 24 hours after i.p. injection of controls or 250 ng of mRNA. (F) Quantification of mCherry MFI across the immune cell compartment and. (G) Experimental layout illustrating the treatment of mice with either scrambled or Lyve1 siRNA. Mice are injected intraperitoneally and after 48 hours peritoneal lavage is harvested for flow cytometry (H-J) and analyzed for %Lyve1 positive macrophages in Tim4+ or Tim4-cells (I) as well as Lyve1 MFI (J). Experimental layout illustrating the treatment of WT or H11Cas9 mice with aTim4-targeted ShredR LNPs i.p. using PBS and WT animal as control (K). Flow cytometric analysis of Tim4+ LPMs in peritoneal lavage of WT (left) and H11Cas9 (right) mice 48 hours after i.p. of anti-Tim4-ShredR LNPs (L) and (M) relative % of Tim4+ cells compared to the vehicle control (PBS). Error bars show ±SEM and statistically significant differences for E, F, I, J was calculated using 2-way ANOVA followed by Turkeys multiple comparison test,, whereas M was calculated using an unpaired t-test. *, P < 0. 05, **, P < 0. 01, ***, P < 0.001, ****, P < 0.0001

As Lyve1 and Tim4 was found to define separate populations by both scRNA-seq and flow cytometry (fig. 4B,C), we initially focused on validating subset-specific targeting to these cells by preparing high PEG SM102-LNPs encapsulating mRNA for mCherry conjugated to anti-mouse Tim4 antibody (αTim4-mCherry(mRNA)-LNPs). Next, subset-specific targeting was evaluated by injecting naïve mice *i*.*p*. with the αTim4-conjugated LNP formulation, and peritoneal cells were harvested for flow cytometric analysis after 24 hours (fig. 4C). The results demonstrated a specific targeting of Tim4+ LPMs when injecting the mice with Tim4-targeted SM102-LNPs, as only Tim4+ LPMs showed a high specific increase in mCherry expression (fig. 4E-F). In contrast, LPMs not expressing Tim4 or mice injected with control formulations only showed minimal mCherry expression in macrophages.

Having successfully validated targeting of Tim4+ cells using the aTim4-LNP formulation, we next sought to establish subset specific knock-down of Lyve1 in Tim4+, Lyve+1 cells only, by encapsulating Lyve1 siRNA in aTim4-LNPs. Following initial titration of siRNA loaded aTim4-LNPs, *in vivo* we found that specific know down of Lyve1 expression in Tim4+ cells was achieved with a single dose of 250 ng siRNA (fig S4B-C). Importantly, increasing the dose to a concentration often used in other *in vivo* LNP studies ^32^ increased Lyve1 knockdown in Tim4-cells, suggesting a non-targeted uptake. To further validate the specific knockdown of Lyve1 in Tim4+ macrophages, we next repeated the siRNA-LNP experiment including an aTim4-LNPs scrambled siRNA control. As before, injecting mice *i*.*p*. with aTim4-Lyve1(siRNA)-LNPs resulted in specific knockdown of Lyve1 expression in Tim4+ cells as observed by the strong reduction of Tim4+, Lyve1+ cells and the concomitant reduction in Lyve1 MFI in the remaining Lyve1+, Tim4+ cells (fig. 4I-J). This was not observed in the mice injected with aTim4-Scrambled(siRNA), with no MFI change relative to PBS.

Lastly, as a final proof of principle and to validate the ability of the targeted SM-102 LNPs to deliver functional sgRNAs for use in CRISPR/Cas9 mediated gene editing *in vivo*, we next encapsulated ShredR sgRNA in the aTim4-LNPs and injected them *i*.*p*. in wildtype C57Bl/6 mice and H11^Cas9^ mice (fig. 4K). These mice express spCas9 across multiple cell types. After 48 hours we evaluated the %of Tim4+ macrophage in the peritoneum and found that the percentage of Tim4 expressing cells was significantly reduced in Cas9 expressing mice (fig 4M).

## Discussion

Extensive research has effectively documented the important functions of macrophages in both tissue homeostasis and disease development and in the context of caner, several therapeutic strategies depleting or blocking macrophage infiltration has been tested in the clinic^33^. Yet, so far these strategies have shown limited effect. However, similar to most other macrophages, tumor-infiltrating macrophages are highly diverse, and whereas a high presence of macrophages is generally linked to a poor prognosis, certain TAM subsets can be linked to positive outcomes, likely due to their key roles in driving protective anti-tumor immune responses^34^. Therefore, selectively targeting the tumor-promoting macrophage subsets or their tumor-promoting functions, while preserving the potential anti-tumor populations, could provide substantial clinical benefits across various cancers.

To this end, there is currently a substantial interest in the macrophage field to identify and characterize macrophage heterogeneity and function of individual macrophage subsets. In many cases, researchers combine single cell transcriptomics with laborious techniques such as cre dependent mouse models or even pan-targeting strategies such as clodronate liposomes which do not readily offer the ability to target macrophage heterogeneity.

Importantly, Cre-based models, such as LyzM-Cre, Csf1r-Cre and Cx3Cr1-Cre have been instrumental for elucidating macrophage function *in vivo* and recently also for providing experimental evidence on macrophage origin^13^. However, in most cases, setting up Cre models require extensive and time-consuming breeding programs. Furthermore, in many models, Cre expression is not fully restricted to macrophages^12^. This is for example the case for one strain of Csf1r-Cre, where Cre activity has been observed in neutrophils, dendritic cells and T cells^35^. In our lab, we have observed issues when using the constitutive Cre model CD163-iCre. Here we observed Cre activity in B cells, most likely as a cause of transient expression of the iCre, during a precursor stage^30^. Though this could potentially be circumvented using the inducible Cre, this would present another potential caveat. Tamoxifen, an often-used inducer, is toxic to work with, especially for pregnant women, but it is also toxic for mice, causing alterations in the macrophage phenotype ^36^, and in the immune cell compartment in general ^17^.

In the present study, we have demonstrated the development of a robust toolbox, enabling efficient delivery of RNA to macrophages. We have shown that the SM-102 LNPs efficiently encapsulates both siRNA, sgRNA and mRNA and effectively target this to macrophages *in vitro* and *in vivo*. Moreover, using antibody targeted LNPs, we were able to achieve subset specific targeting of distinct macrophage populations *in vivo* and provide the first proof-of-principle for a LNP based approach for subset specific depletion or knockdown of protein expression, offering an alternative to more non-specific or complicated conventional approaches.

One of the major benefits of the LNP model is the flexibility as it provides a platform where any RNA modulator can be loaded and specifically delivered to macrophages *in vitro* and *in vivo*, allowing investigators to screen large libraries of targets. Using the SM-102 LNPs, the RNA cargo can be readily exchanged for RNA targeting another signaling pathway, with quick turnover available for new strategies. By knocking down or overexpressing large numbers of genes in parallel, researchers can rapidly identify genes involved in key cellular processes or identify potential therapeutic targets. Additionally, a major advantage of the LNP model is apparent when using the targeted approach, as developing new targeting particles only requires an available antibody and the specific functional RNA molecule which is often commercially available. This is in stark contrast to setting up new CRE based models that often require much more work, such as rederivation and lengthy breeding strategies.

In recent years, a number of new genome editing technologies have emerged with the CRISPR-Cas9 system being the most used. To explore this system in the LNP based approach we generated the murine ShredR sgRNA that targets short interspersed nuclear elements (SINEs) and show that this sgRNA, when combined with Cas9 expression can drive macrophage depletion. In this study, we used two different Cas9 mouse strains that either expressed the Cas9 enzyme ubiquitously (Cas9-H11) or macrophage restricted (Csf1r^Cas9^). In both cases we observed efficient macrophage depletion, highlighting the high degree of flexibility and specificity of the system, as LNP-mediated delivery of sgRNA may be combined with Cre-specific expression of floxed transgene Cas9 enzyme to allow for a combination of genomic and proteomic targeting of individual macrophage subsets. Moreover, Cas9 enzyme may also be delivered to macrophages as mRNA, allowing for a dual protein targeted approach.

In addition to the SM102 LNPs presented here, several groups have recently published methods to transfect macrophages with RNA using LNPs. One example is the recent paper by Huang et al (2022), where the authors were able to achieve successful transfection of macrophages, using surfactant-derived LNPs *in vitro*^*37*^. However, the transfection efficiency using this LNP formulation was less than observed in HEK cells. In a different study, recently published by Zhao et al (2024)^32^, the authors used LNPs coated with antibodies, to deliver siRNA-LNPs intra-nasally to provide proof-of-concept for *in vivo* delivery. While this study used a similar method, as described here, the authors focused on pan-targeting of macrophages by using a F4/80 antibody. Even though this study validates the ability to deliver RNA to macrophages *in vivo*, the authors also observed a relatively low transfection efficiency compared to the data presented here (less than 50%) which was accompanied by a substantial level of unspecific uptake of their LNPs in CD68 negative cells, although the exact level was not quantified. To achieve this level of efficiency, the authors used a dose of 1.25 mg/kg, which is equal to approximately 25 μg siRNA per 20g mouse, which is approximately 100 times higher compared to the dose used in this study. The high RNA dose resulted in excessive immune infiltration which had to be inhibited with an injection of dexamethasone 1 hour prior to LNP inoculation. Importantly, although this has not been studied in detail in the study presented here, our data using *i*.*p*. injection and doses at 1.5 μg per mouse or less, do not indicate any immune reaction nor infiltration of inflammatory immune cells.

In conclusion, in the present study we have presented a readily available, versatile toolbox that enables subset specific editing of cell functions using readily available RNA technologies such as siRNA or CRISPR-Cas9. While this is not a replacement for the well characterized systems such as the Cre-LoxP system, this method is a relatively low cost, time saving alternative, allowing you to examine multiple signaling pathways, with no effect on viability of the cells.

## Limitations of study

In the present study we have focused on proving *in vivo* use of the LNP formulation using *i*.*p*. injection only and although we believe the formulation is widely available, other injection routes will need to be optimized separately. Despite our efforts to reduce non-specific uptake when using the Tim4 targeted LNPs a low degree of mCherry expression was seen in macrophages not expressing Tim4, although with translating into a functional KD when encapsulating siRNA. This unspecific uptake likely reflects the saturation of Tim4 receptors at higher doses and future work should focus on optimizing RNA-LNP dosage and improving targeting specificity to minimize off-target effects.

## Acknowledgements

We thank Martin Kristian Thomsen, Felicity Davies and Joanna Kalucka (Department of Biomedicine, Aarhus, Denmark) for Cas9, Csf1r-Cre and Cas9-EGFP^Lox-Stop-Lox^ mice and Gitte Fynbo Biller for excellent technical assistance. We also thank Ronni Nielsen and the Functional Genomics & Metabolism unit (SDU, Odense, Denmark) for help with RNA sequencing. These studies were supported by a grant to AE from Aarhus University Research Fund, the Novo Nordisk Foundation (NNF20OC0065510 and NNF22OC0080192) and the Danish Cancer Society (R302-A17596) as well as a grant to MNA from the Novo Nordisk Foundation (NNF22OC0079892). Flow cytometry and cell sorting was performed at the FACS Core Facility, Aarhus University, Denmark. The 5-laser ID7000 is a generous gift from the Novo Nordisk Foundation, grant number NNF210C0066798.

## Data availability

Single cell RNA-seq data is available upon reasonable request.

## Author contributions

AE and RKR conceptualized the study and designed the experiments. RKR developed LNP formulations and RKR + HM developed the pipeline for antibody-functionalization. HM and MWK performed Incucyte imaging with support from MNA, qPCR was performed by IHM, *In vivo* studies were performed by RKR, JMG and AE. JMG performed sorting and library prep for scRNAseq analysis, and LMS performed scRNA sequencing dataanalysis. RKR performed the *in vitro* ShredR experiment and RKR and MBT developed the sgRNA-ShredR platform. RKR and AE wrote the manuscript, and all other authors commented.

