## Supplemental figures for "Targeted delivery of RNA-based therapeutics enables functional analysis of macrophage subpopulations"

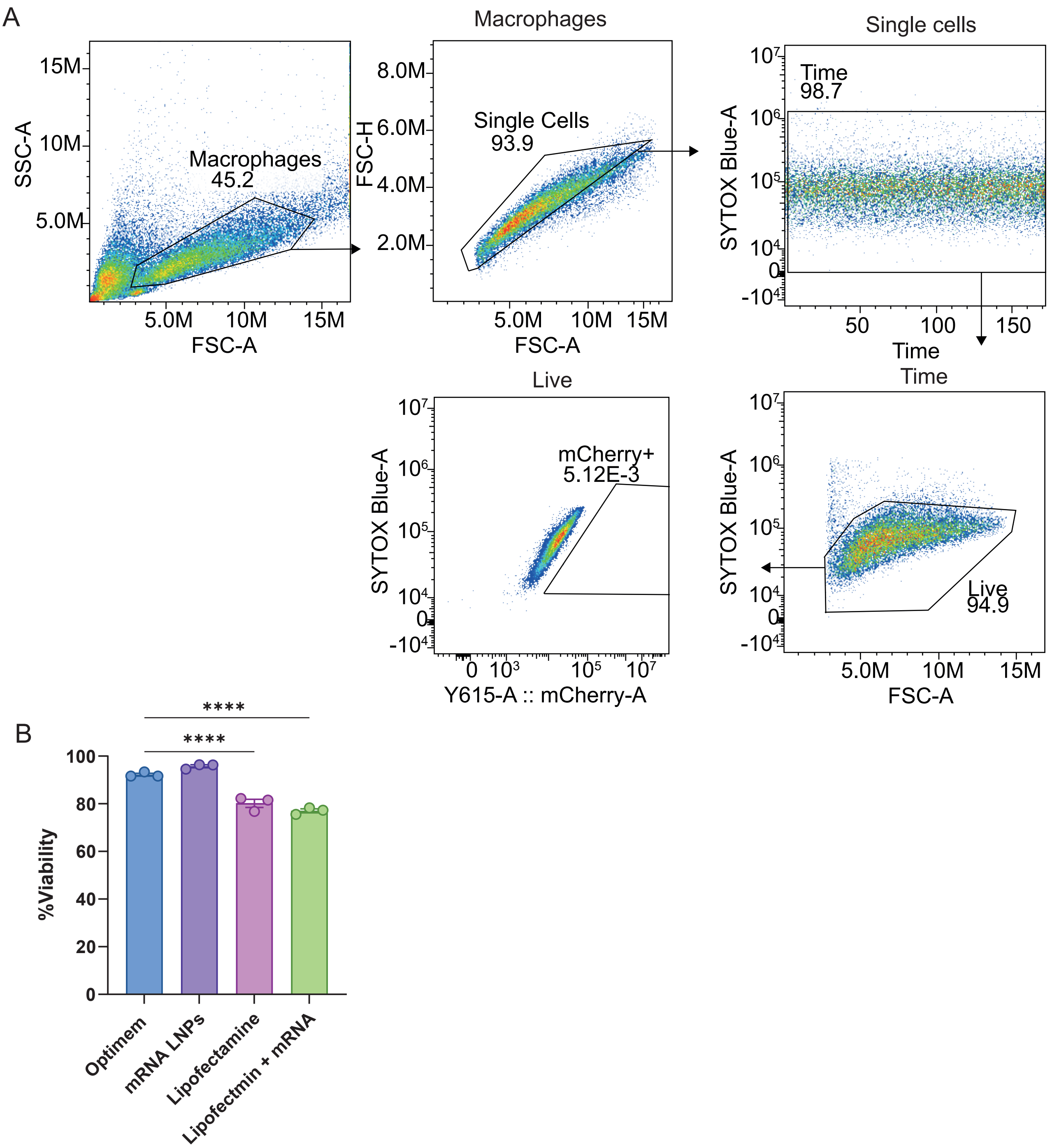


**Figure S1 - Gating strategy and viability for *in vitro* characterization of SM-102 LNPs** A -Gating strategy for BMDMs. B - Viability of BMDMs after treatment with media alone, mRNA-LNPs or lipofectamine alone, or lipofectamine with mRNA. Error bars show± SEM, statistics: One-way ANOVA with Dunnetts multiple comparisons test. ****, P < 0.0001

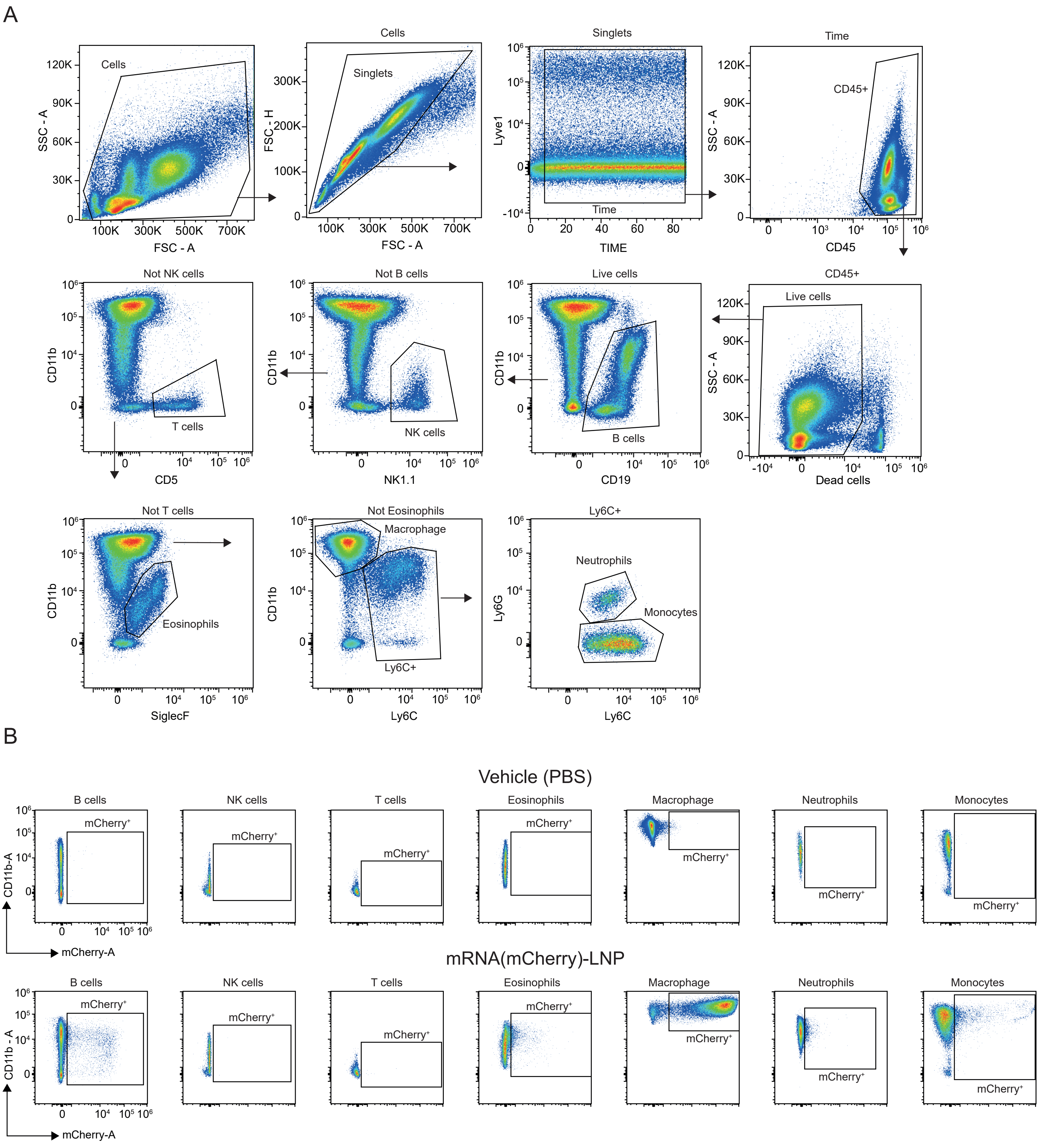


**Figure S2 - Gating strategy for *in vivo* characterization of SM-102 LNPs** A -Gating strategy for all cell types B - Example gates for mCherry positive cells in PBS or mCherry LNP treated mice

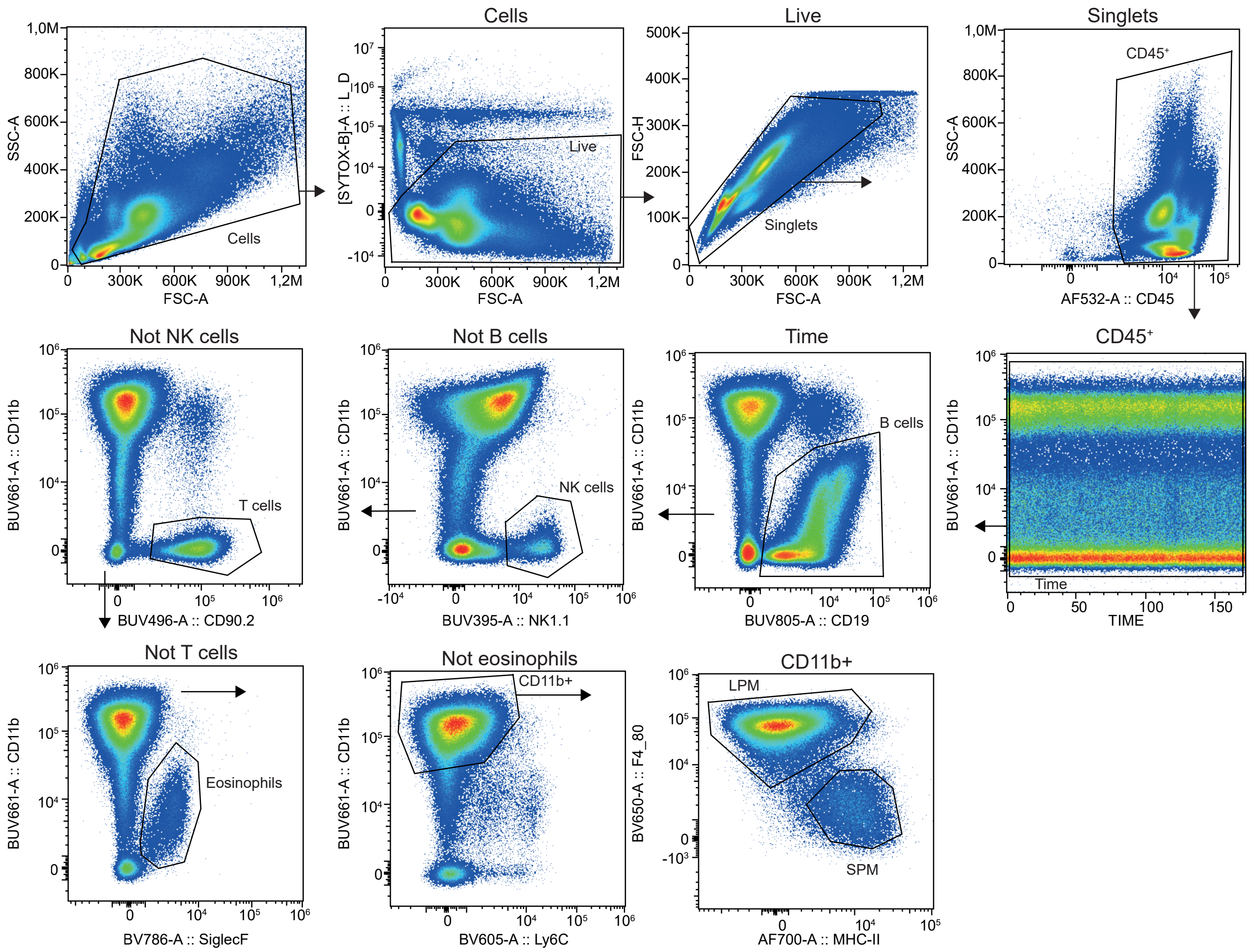


**Figure S3 - Gating strategy for the targeted SM-102 mRNA LNPs** Gating strategy for all cell populations.

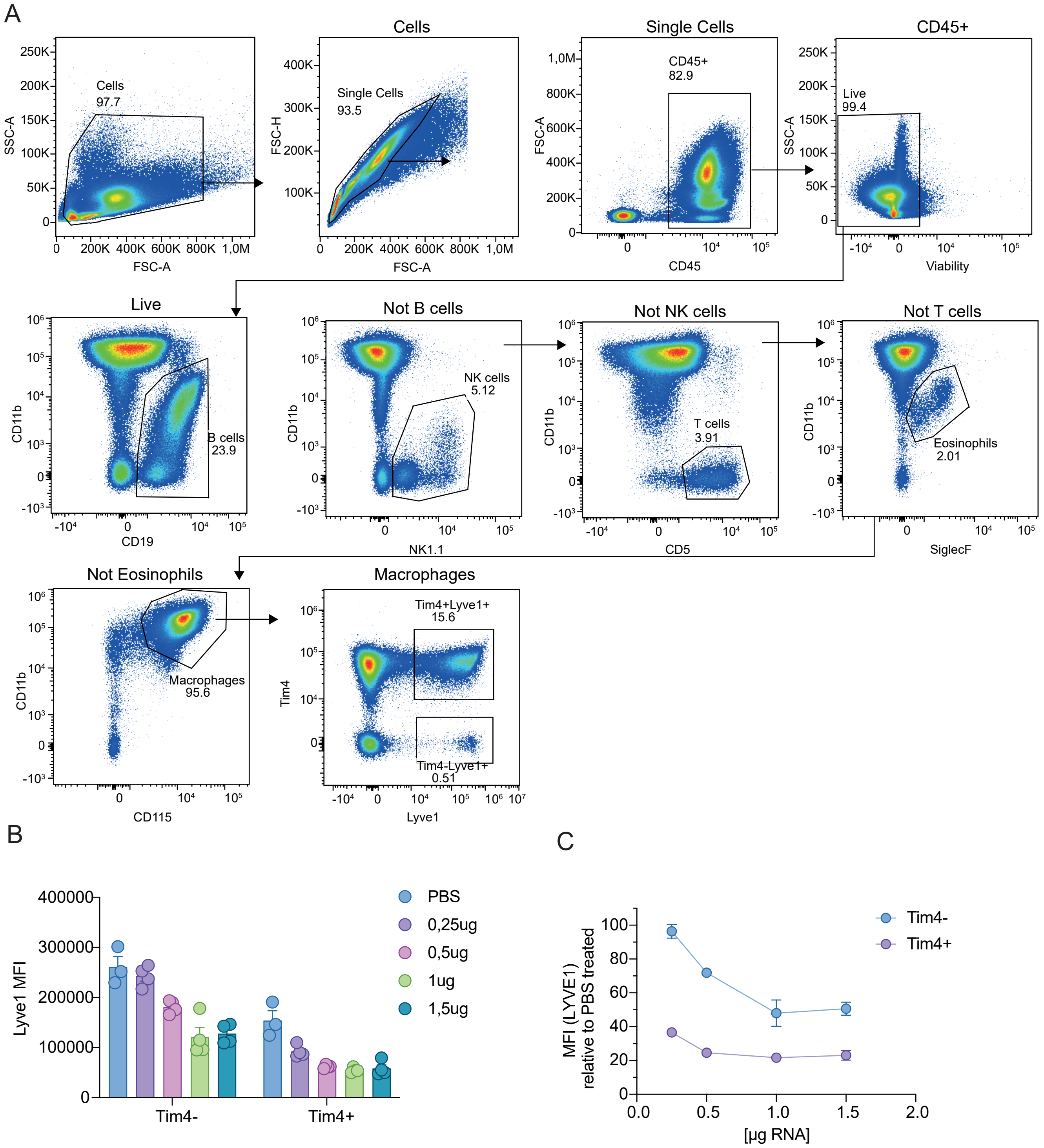


**Figure S4 - Gating strategy for the targeted SM-102 siRNA LNPs** A - Gating strategy. B - Lyve1 MFI of the Tim4+, Lyve1+ macs and the Tim4-. Lyve1+ macs. C - MFI of Lyve1 relative to vehicle treated mice (PBS). Error bars show ± SEM

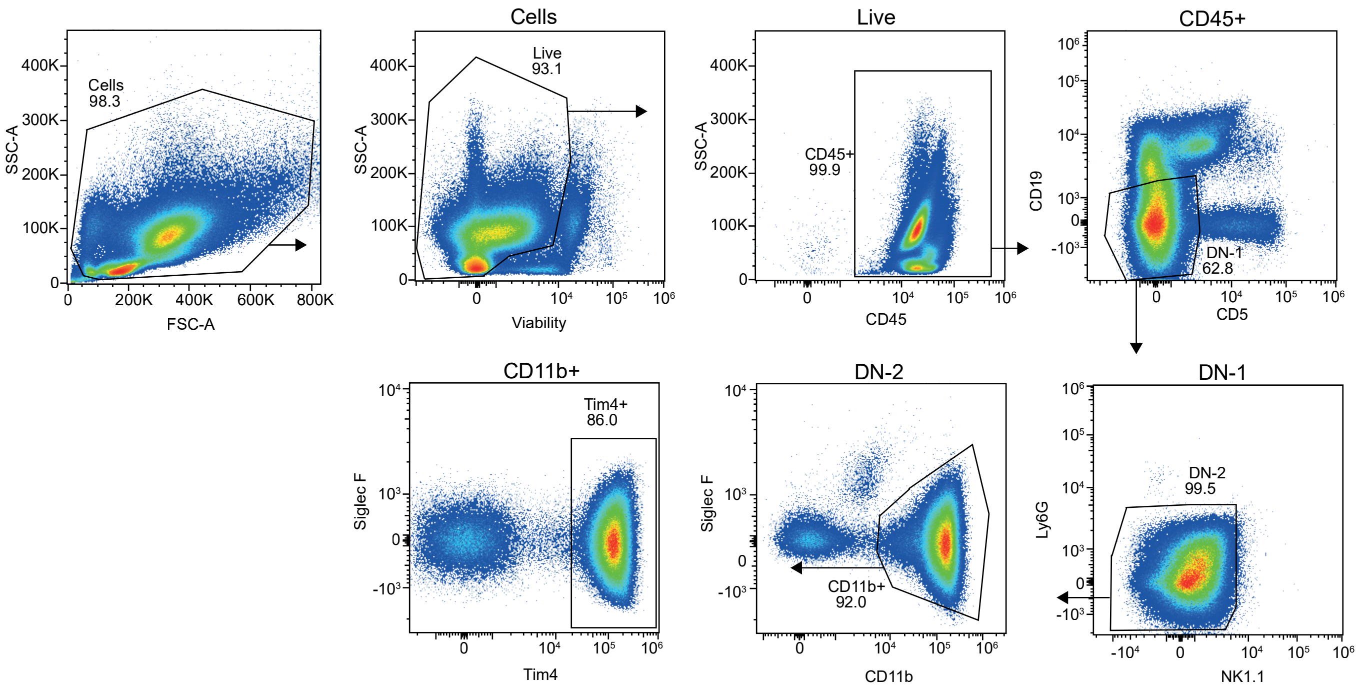


**Figure S5 - Gating strategy for the targeted SM-102 sgRNA LNPs** Gating strategy for Tim4+ myeloid cells.
